# Extended haplotype phasing of *de novo* genome assemblies with FALCON-Phase

**DOI:** 10.1101/327064

**Authors:** Zev N. Kronenberg, Arang Rhie, Sergey Koren, Gregory T. Concepcion, Paul Peluso, Katherine M. Munson, Stefan Hiendleder, Olivier Fedrigo, Erich D. Jarvis, Adam M. Phillippy, Evan E. Eichler, John L. Williams, Tim P.L. Smith, Richard J. Hall, Shawn T. Sullivan, Sarah B. Kingan

## Abstract

Haplotype-resolved genome assemblies are important for understanding how combinations of variants impact phenotypes. These assemblies can be created in various ways, such as use of tissues that contain single-haplotype (haploid) genomes, or by co-sequencing of parental genomes, but these approaches can be impractical in many situations. We present FALCON-Phase, which integrates long-read sequencing data and ultra-long-range Hi-C chromatin interaction data of a diploid individual to create high-quality, phased diploid genome assemblies. The method was evaluated by application to three datasets, including human, cattle, and zebra finch, for which high-quality, fully haplotype resolved assemblies were available for benchmarking. Phasing algorithm accuracy was affected by heterozygosity of the individual sequenced, with higher accuracy for cattle and zebra finch (>97%) compared to human (82%). In addition, scaffolding with the same Hi-C chromatin contact data resulted in phased chromosome-scale scaffolds.

## INTRODUCTION

A high-quality reference genome is an indispensable resource for basic and applied research in biology, genomics, agriculture, medicine, and many other fields^1–3^. Technological innovations in DNA sequencing, long-range genotyping, and assembly algorithms have led to rapidly declining costs of sequencing and computation for genome assembly projects^4^. As researchers are now able to pursue genome projects for outbred, non-model, diploid and polyploid organisms, a major challenge in *de novo* assembly is accurate haplotype resolution. Most genome assemblers “collapse” multiple haplotypes into a single consensus sequence to generate a pseudo haploid reference. Unfortunately, this process results in mosaic haplotypes with erroneously associated variants not present in either haplotype and concomitant effects on biological inference^5–7^.

Three approaches to haplotype resolution in long-read diploid genome assembly have been described. The most recently reported, trio binning, depends on short-read data of the parents to identify parent-specific *k*-mers, which are then used to separate long reads from the offspring prior to assembly into maternal and paternal bins^8^. These parent-specific bins are then assembled into two haploid references, thus entirely avoiding the added complexity of diploid assembly. This method provides extremely accurate phased assemblies representing each parental contribution, but requires that samples of the parents are available, which may not always be the case for samples of interest. A second approach phases reads by mapping to an existing reference genome to infer haplotypes, followed by long-read partitioning and assembly^9–12^. Read-based phasing methods require that a reference assembly is available, and depend on single nucleotide variant (SNV) calling that has associated error. The third approach is to separate haplotypes during the genome assembly process as implemented by FALCON-Unzip for long reads^13^ and Supernova for short reads^14^. FALCON-Unzip outputs two types of genomic contigs: primary contigs, which are highly contiguous pseudo-haplotypes containing both phased and unphased haplotypes, and haplotigs, which are shorter phased contigs that represent the alternate alleles in heterozygous regions of the primary contig. The length of the phase blocks (haplotig plus homologous primary contig region, see Fig. 1) produced by FALCON-Unzip are limited by the distribution of read length, depth of coverage, and the degree and distribution of heterozygosity in the diploid genome. Regions of low heterozygosity are collapsed into non-haplotype-resolved sequence because they contain insufficient information for read phasing.

**Figure 1.**
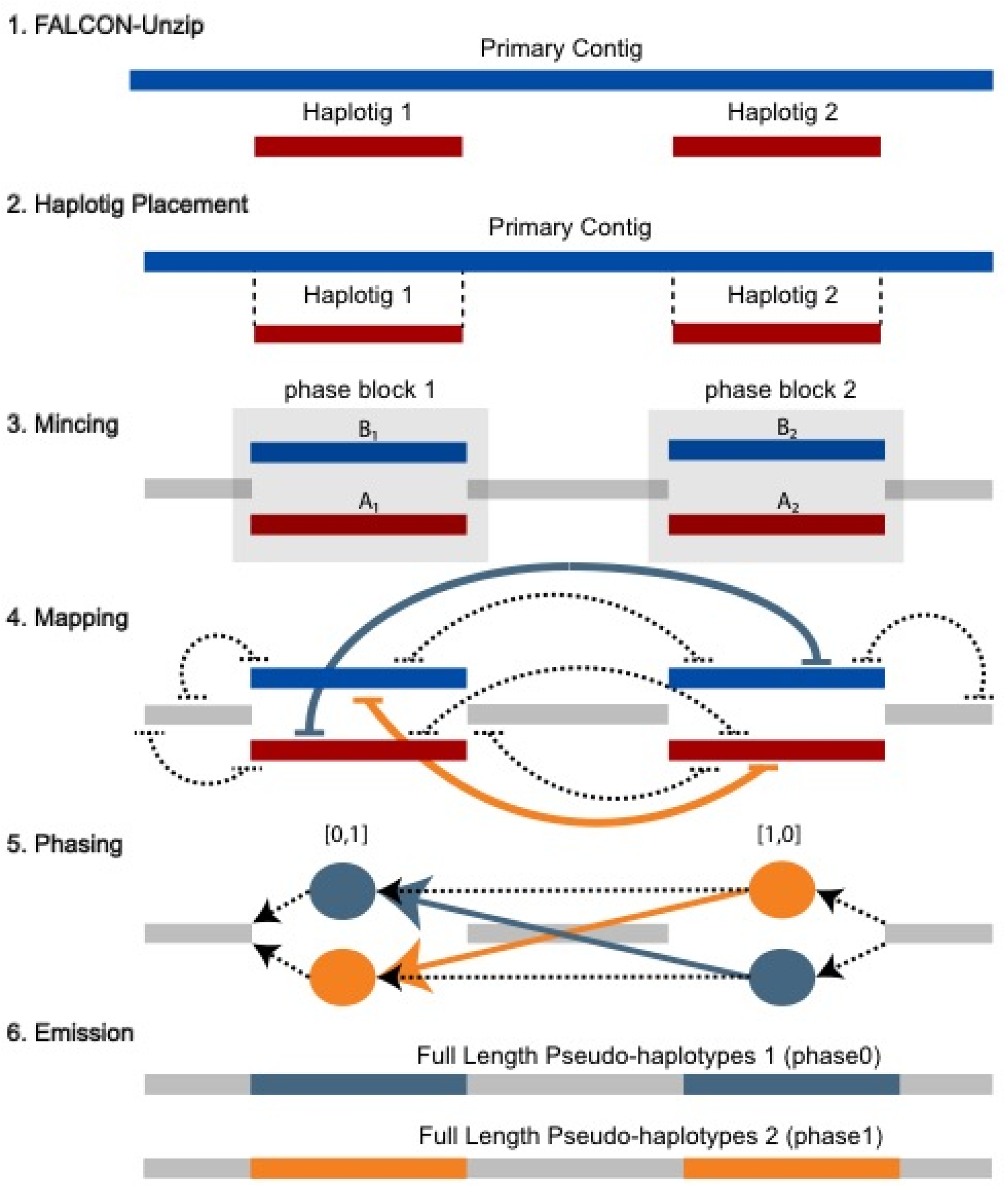
Overview of FALCON-Phase method. 1. FALCON-Unzip assembly consists of long primary contigs (blue) and shorter alternate haplotigs (red). The region where a haplotig overlaps a primary contig is a “phase block” and is referred to as being “unzipped” because two haplotypes are resolved. Regions of the primary contig without associated haplotigs are referred to as “collapsed” because the haplotypes have low or no heterozygosity. 2. A haplotig placement file specifies primary contig coordinates where the haplotigs align. 3. This placement file is used to “mince” the primary contigs at the haplotig alignment start and end coordinates. Mincing defines the “phase blocks” (A-B haplotype pairs, blue and red) and collapsed haplotypes (grey). 4. Hi-C pairs are mapped to the minced contigs and alignments are filtered. 5. Phase blocks are assigned to state 0 or 1 using the phasing algorithm. 6. The output of FALCON-Phase is two full length parental pseudo-haplotypes for phase 0 and 1. These sequences are of similar length to the original FALCON-Unzip primary contig and the unzipped haplotypes are in phase with each other.

To eliminate the haplotype switch errors in FALCON-Unzip primary contigs, we developed FALCON-Phase. It addresses the problem of phase switching between FALCON-Unzip phase blocks by integrating ultra-long-range genotype (>1Mb) information from Hi-C read pairs^15^. Unlike read-based phasing methods, FALCON-Phase does not require an existing reference genome or SNV calling, but instead relies on the haplotype-specific mapping pattern of informative Hi-C read pairs between long-read generated contigs of the same individual^9,12,16^. In contrast to trio binning, FALCON-Phase is based entirely on sequence of the individual being assembled, and does not require parental samples or sequence.

## RESULTS

### FALCON-Unzip Assembly and Processing

We generated contig draft *de novo* assemblies for three vertebrate species (human, zebra finch, and cattle) using FALCON-Unzip (see Table S1 for raw assembly statistics, Table S2 for data availability). We curated the initial assemblies to break chimeric contigs manually (see methods) and used purge haplotigs^17^ to remove duplicated haplotypes from the primary contig set. This ensured that the starting primary assembly was haploid and that alternate haplotig sequences were each associated with a primary contig. The primary assemblies ranged from ~1-3Gb in size (contig N50 length of ~4-30 Mb) with 84-88 % of the genomes haplotype-resolved. Average haplotig length, which is equivalent to average phase block size, ranged from 188-452 kb (Table 1). We then applied FALCON-Phase to these assemblies, which first defines phase blocks by aligning the alternate haplotigs to their associated primary contigs, then uses mapping density of Hi-C read pairs to bin haplotype sequences that are on the same chromosome, and finally expands the homozygous sequences into both pseudo-haplotypes (Figure 1). Users can specify “pseudo haplotype” or “unzip” output formats, the former having the same “collapsed” sequence in both pseudo-haplotypes, the latter matching the FALCON-Unzip assembly format.

**Table 1.** Summary statistics of inputs: starting genome assemblies and Hi-C data

| Sample | Zebra Finch | Cattle | Human |
| --- | --- | --- | --- |
| Heterozygosity (%) | 1.57 – 1.72 | 0.65 - 0.93 | 0.17 – 0.21 |
| Primary Assembly Length (Gb) | 1.05 | 2.71 | 2.89 |
| Primary Contig N50 (Mb) | 3.48 | 31.4 | 26.3 |
| Mean Phase Block Length (Mb) | 0.188 | 0.452 | 0.312 |
| Proportion of Genome Phased (%) | 87.6 | 87.7 | 84.0 |
| Average number of Hi-C links between phase blocks on the same primary contig (pre/post) filtering | 92.5 / 31.5 | 20.39 / 4.79 | 44.79 / 2.42 |

FALCON-Phase aligns Hi-C read pairs to both the collapsed regions and phase blocks using bwa^18^. We found that most reads (56-88%) mapping to the assembly did not contain haplotype-specific variants, and therefore had low or no phasing information (Table S3). By requiring both Hi-C read pairs to have a map quality greater than 10, we obtain a haplotype-specific set of Hi-C reads. In the cattle assembly, for example, 16.3% of the Hi-C read pairs pass this filtering (64.5 M / 395 M), resulting in 23.8 Hi-C read pairs per kilobase (kb) across the genome (Table S3). A matrix is then generated from the counts of retained Hi-C read pairs mapping between phase blocks.

### Contig-Phasing Performance

We found that phasing performance was impacted by the level of heterozygosity (Table 2). The zebra finch assembly has the highest heterozygosity (~1.6%; Table 1) and 98.3% of the phase blocks were properly phased (Table 2); the cattle sample has a lower heterozygosity (~0.9%), with a slightly lower 97.3% of the phase blocks properly phased; human has a ~ 4 to 9-fold lower rate of heterozygosity (~0.2%) than the other two species, and FALCON-Phase achieved only 81.8% accuracy. These findings suggest that lower rates of heterozygosity, where there are fewer informative Hi-C reads after map-quality filtering, interferes with the ability to phase. Consistent with this idea, we observed the fewest Hi-C links between phase blocks in human after filtering (Table 1; Figure S1). A potential alternative explanation is that the choice of restriction enzyme in the Hi-C library construction affects genome coverage and, therefore, ability to phase. This is consistent with the data in that the human Hi-C library was constructed with a single enzyme having a six-base recognition sequence (HindIII), the cattle library was constructed with a single enzyme but four-base recognition sequence (Sau3AI), and the zebra finch library was constructed using multiple enzymes. There is an ongoing effort to resolve which Hi-C library preparation is best for human; simulation studies were inconclusive.

**Table 2.** Phasing accuracy

| Sample | Zebra Finch | Cattle | Human |
| --- | --- | --- | --- |
| FALCON-Phase Accuracy* (%) | 98.3 | 97.3 | 81.8 |
| FALCON-Unzip Primary Contig Accuracy (%) | 70.8 | 71.0 | 61.0 |
| FALCON-Unzip Haplotig Accuracy (%) | 94.9 | 98.7 | 96.2 |
| FALCON-Phase Pseudo-haplotype Accuracy (%) | 91.2 | 96.0 | 80.3 |
| Unphased Scaffold Accuracy (%) | 64.1 | 77.8 | 62.9 |
| FALCON-Phase Scaffold Accuracy (%) | 88.4 | 92.4 | 73.9 |
| Trio binned Canu Contig Accuracy (%) | 99.4 | 99.4 | 99.5 |

\* Based on alignment of phase blocks to trio binned Canu parental assemblies. Other measures are based on parental *k-mer* counting.

Using the phase assignment for the pair of haplotypes in each phase block, we generated two pseudo-haplotypes, phase0 and phase1. These pseudo-haplotypes are similar to the primary contigs of FALCON-Unzip except that the phase blocks should originate from the same parent rather than consisting of a mix of maternal and paternal haplotypes. We assessed the overall phase accuracy of the primary contigs and haplotigs from FALCON-Unzip and the resulting pseudo-haplotypes after FALCON-Phase by counting parental *k-mers* identified in Illumina data from the parents. This stringent measure penalizes every *k-mer* that is contained within a phasing switching error. For all three genomes, the FALCON-Unzip haplotigs already had phasing accuracy greater than 95% (Table 2, see also^19^) whereas the primary contig phasing accuracy ranged from 71% for zebra finch and cattle to 61% for human. After applying FALCON-Phase, the resulting pseudo-haplotypes had an increase in the proportion of markers from one parent: correct phasing in the zebra finch and cattle increased to 91% and 96%, respectively, and in human, to 81.8% (Figure 2, Table 2). While each pseudo-haplotype contig contained a majority of markers from one parent, the set of phase0 or phase1 contigs are a mix of maternal and paternal pseudo-haplotypes (Figure 2d-f). As a ceiling control, *k-mer* analysis of trio binned Canu assemblies have >99% phasing accuracy for these genomes. We also evaluated the phase accuracy for a supernova assembly of the human sample and determined it to be 74% for both pseudo-haplotypes (Figure S2).

**Figure 2.**
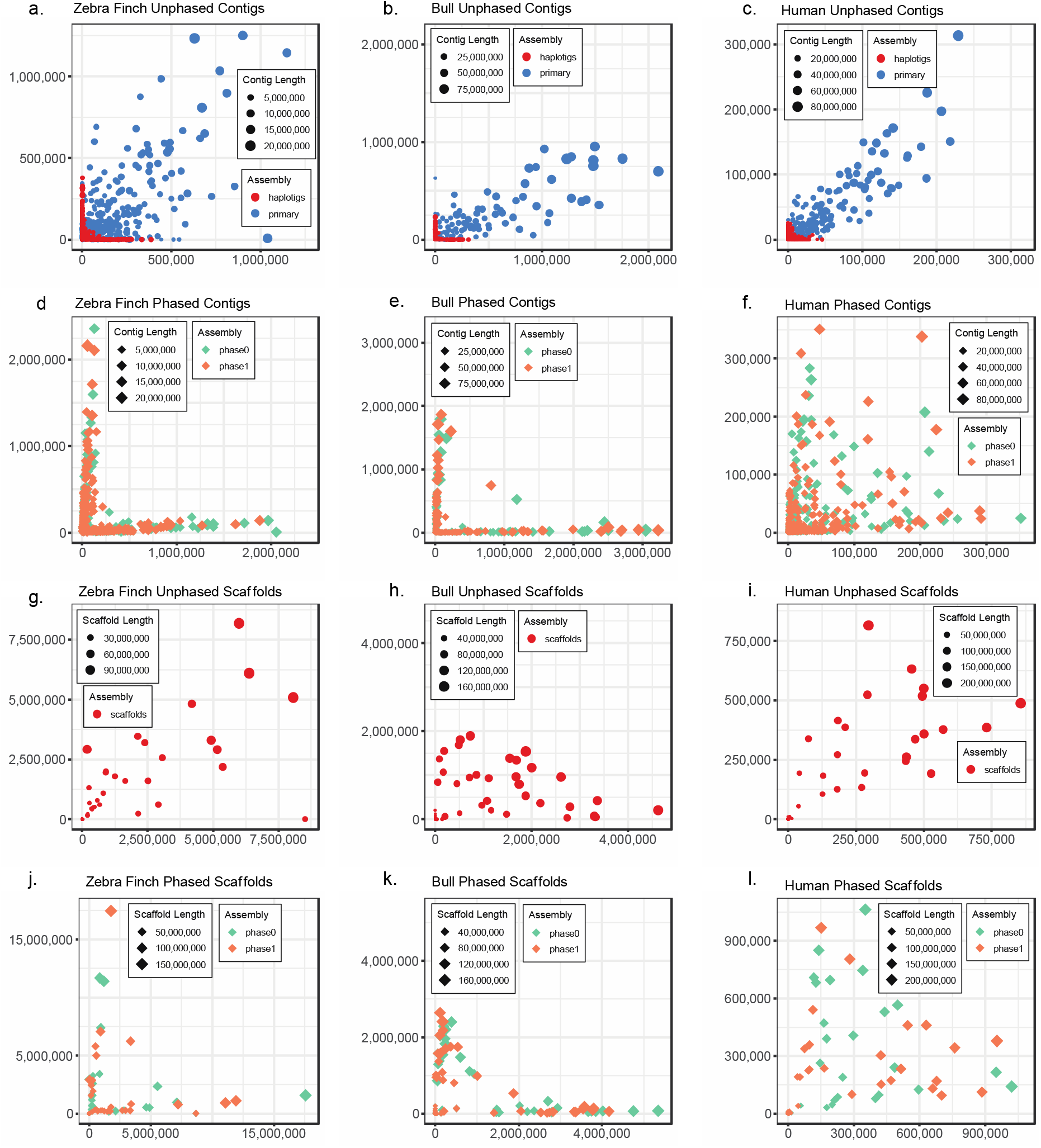
Parental k-mer counts for contigs and scaffolds before and after phasing. Markers from mother are on the X axis, father on the Y-axis. Contig or scaffold size are indicated by size of the data point. Tested species are arranged in order of decreasing heterozygosity (zebra finch: A,D,G,J; cattle: B,E,H,K; human: C,F,I,L) A-C. Unphased FALCON-Unzip contigs. Primary contigs (blue) are large but contain a mixture of markers from mother and father. Haplotigs contain less of a mixture of parental markers, but are smaller. D-F Contigs after phasing. Phase0 and Phase1 contigs are of similar length to the FALCON-Unzip primary contigs and have less mixing of parental markers within contigs. G-I, Unphased scaffolds, phase0 contigs only. J-L, Phased scaffolds contain less mixing of markers than unphased scaffolds. As with the contigs, the phase0 and phase1 scaffolds sets contain maternal and paternal haplotypes.

### Scaffold Phasing

Contigs from phase0 pseudo-haplotypes were scaffolded into chromosome-scale sequences using Proximo Hi-C (Phase Genomics, Table S5). A second round of phasing was performed on the scaffolds using FALCON-Phase and performance was evaluated using parental *k-mer* counts in the unphased versus phased scaffolds (Table 2). For cattle and zebra finch, overall parental scaffold phasing accuracy was 92.4% and 88.4%, respectively, and in the human, 73.9% of their markers were from one parent after the second round of phasing. Thus, FALCON-Phase is the only method which generates chromosome-scale phased assemblies from a diploid genome without requiring parental information. Additional information is necessary to compile a maternal or paternal set of scaffolds however, as the phase0 and phase1 sets contain a mix of the parental sequences.

## DISCUSSION

The ultimate goal of genome assembly is to faithfully represent each chromosome in the organism from telomere-to-telomere. To do so, assembly methods must account for sequence similarity between homologous maternal and paternal chromosomes, in order to prevent collapsed haplotypes, which may result in incomplete or erroneous representations of the underlying biological sequence^7,8^. The FALCON-Unzip genome assembler is able to identify heterozygous regions of a genome as bubbles in assembly graphs, and “unzip” those bubbles further by phasing and reassembling reads using single nucleotide variants^13^. However, FALCON-Unzip cannot phase entire primary contigs. To address this limitation, we designed FALCON-Phase, which uses Hi-C data to extend the phase blocks to the contig and scaffold scales. Here, we have demonstrated that FALCON-Phase improves accuracy for heterozygous diploid genome assemblies, without parental or population sequence data.

FALCON-Phase, in conjunction with FALCON-Unzip, is thus an attractive method for generating high quality reference genomes of samples with moderate to high heterozygosity for which parents are not available. This approach may be useful for large-scale genome initiatives which source samples of diverse origins, including invertebrate disease vectors, agricultural pests, or other wild-caught individuals. The method utilizes two technologies common in generating highly contiguous genome assemblies: PacBio continuous long reads (CLR) and Hi-C. While Hi-C has recently been used for scaffolding^20,21^, we find that the same data can also be used for contig or scaffold phasing. The accuracy of phasing increases with higher heterozygosity; the human sample had the lowest heterozygosity, and consequently fewer haplotype-informative Hi-C links. However, the comparatively low heterozygosity levels encountered in human are notably rare among animals, in particular wild populations. FALCON-Phase is thus expected to be valuable for population and conservation genomics^22,23^.

FALCON-Phase relies on a diploid assembly that is curated as a haploid set of primary contigs plus alternate haplotigs that are each assigned to a primary contig. Any primary contig is treated as if it were diploid and will be duplicated in the pseudo-haplotype output. Contigs from hemizygous regions of the genome like the sex chromosomes or mitochondrial sequences should not have phase-switch errors and may be removed prior to running FALCON-Phase. Similarly, we recommend removing duplicated haplotypes in the primary assembly manually or with automated tools such as purge haplotigs^17^. Lastly, additional curation to remove chimeric contigs that join unlinked loci is recommended prior to application of our method^21^. While curation is optional, it is recommended as FALCON-Phase does not quality control the input assembly.

The phasing algorithm at the core of FALCON-Phase could be adapted to utilize other long-range contact data types. The input matrix is simply a count of contacts between all pairs of sequences in an assembly. Instead of Hi-C data, BAC-end sequences, read clouds, or optical maps could be transformed into the required input for FALCON-Phase. Hi-C was chosen over the other technologies because it provides ultra-range contact information (>1Mb) which enables chromosome-scale phase blocks. Similarly, the input sequences could consist of phase blocks derived from methods other than a FALCON-Unzip assembly, such as phase blocks generated through resequencing, variant calling, or pseudo-haplotypes generated from other long-read combinations^24^. The simple, yet novel, concept of skirting variant calling reduces the number of steps and overall runtime of phasing pseudo-diploid assemblies. For these reasons, we believe FALCON-Phase will be an important algorithmic contribution to the goal of diploid genome assembly.

## MATERIAL AND METHODS

FALCON-Phase has three stages: 1) processing FALCON-Unzip contigs and Hi-C data; 2) application of the phasing algorithm; and 3) emission of phased pseudo-haplotypes (Fig. 1). We implemented FALCON-Phase using the Snakemake language to provide flexibility and pipeline robustness^25^. The pipeline can be run interactively, on a single computer, or submitted to a cluster job scheduler. The code is open source under a Clear BSD plus attribution license and is available through github (https://github.com/phasegenomics/FALCON-Phase).

### FALCON-Phase Method

#### Identification of Phase Blocks

In stage one, the FALCON-Unzip assembly is processed to identify phase blocks: regions of the genome that have been “unzipped” into a maternal and paternal pair of haplotypes. FALCON-Unzip generates contiguous primary contigs representing pseudo-haplotypes and shorter phased alternate haplotigs. A haplotig placement file is generated in the pairwise alignment format^26^ that specifies the alignment location of each haplotig on the primary contig (Fig. 1). Briefly, haplotigs are aligned, filtered, and processed with three utilities of the mummer v4 package: *nucmer, delta-filter* and *show-coords*^27^. Sub-alignments for each haplotig are chained in one dimension to find the optimal start and end of the placement using the *coords2hp.py* script. Finally, non-unique haplotig mappings and those fully contained by other haplotigs are removed with *filt_hp.py*.

The haplotig placement file is used to generate three “minced” FASTA files (Fig. 1), “A_haplotigs.fasta”, “B_haplotigs.fasta”, and “collapsed_haplotypes.fasta”. The “A” haplotigs are the original FALCON-Unzip haplotigs (red in Fig. 1), the “B” haplotigs are the corresponding homologous region of the primary contigs (the alternate haplotype, blue in Fig. 1, part 3-4), and the “collapsed” haplotypes are the unphased or collapsed regions of the assembly (grey in Fig. 1). The pairing of the A and B minced haplotigs in the phase blocks and their order along the primary contig is summarized in an index file, “ov_index.txt”, generated by *primary_contig_index.pl*.

#### Hi-C read mapping

The Hi-C reads are mapped to the minced contigs using *bwa-mem*, with the Hi-C option (−5) enabled^18^. The mapped reads are streamed to *samtools*, removing unmapped, secondary, and supplementary alignments (samtools -F 2316) ^28^. This operation ensures that each mate-pair only contains two alignment records. In the last step of read processing, a map quality score filter of Q10 (for both reads) is applied, removing reads without haplotype-specific sequence. Additionally, we set an edit distance from the reference of less than 5 for both reads. Both more stringent (60) and relaxed (0) map quality filtering resulted in lower phasing accuracy.

#### Contact matrix

The Hi-C mate-pair counts between minced contigs are enumerated into a contact matrix, *M*. Each element, *M_ij_*, in the matrix is later normalized by the number of restriction enzyme sites, *z*, in both the *i^th^* and *j^th^* minced contigs as shown in Equation 1. The raw count matrix is encoded into a binary matrix format.

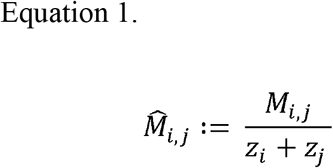

#### Phasing procedure

We designed an algorithm to extend phasing between FALCON-Unzip phase blocks based on Hi-C read pair mapping. The algorithm searches for the optimum set of phase block configurations along a primary contig using a stochastic model. The algorithm is given a list, *C*, of tuples for the phase blocks and their sequential ordering along each primary contig. During initialization, each member of the phase block, except the first, is randomly assigned one of the two possible phase configurations for a diploid organism ∈({[0,1],[1,0]}). The phase assignment is stored in array *T* where 0 corresponds to phase configuration [0,1]. The first phase block along the primary contig is always assigned to the phase configuration [0,1] as its orientation is arbitrary. By fixing the first phase block the search the results are comparable across iterations. Phase blocks are only randomly initialized once before the search begins. The algorithm sweeps along the phase blocks of each primary contig, assigning a phase for the blocks, conditioned on the phase assignment of all previous phase blocks and the Hi-C links between them. The *phaseFreq* function (Equation 2) calculates the frequency of Hi-C links from the current region, *i*, to all past regions, *j*, that have the same phase, i.e. *T_i_* = *T_j_* = 1 = [1,0].

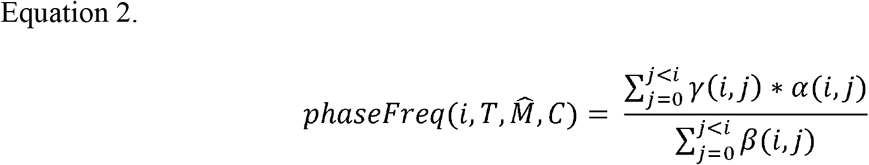

The *phaseFreq* function takes the index of the current phase block, *i*, the phase assignment of all regions associated with a given primary contig, array *T*, the normalized Hi-C count matrix, 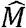, and the *C* array of the phase block tuples. The gamma function (Equation 3) determines if two phase blocks have the same phase assignment, *T*, and if so returns 1. The alpha function (Equation 4) gives the normalized cis counts of Hi-C links between a pair of phase blocks whereas the beta function (Equation 5) returns both the cis and trans counts which is a normalizing constant.

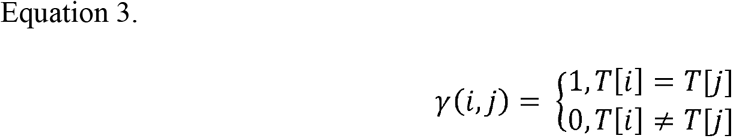

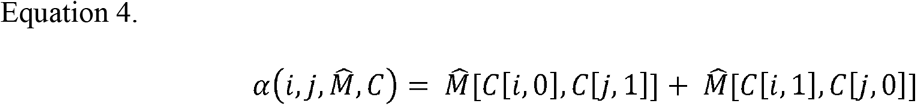

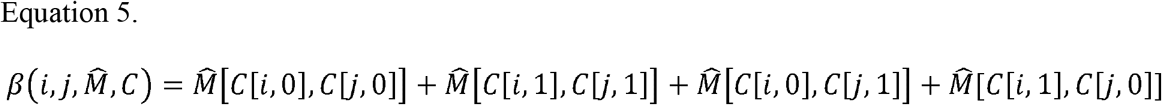

The process of phase assignment across a primary contig is iterated for a burn-in period followed by a scoring period (See Algorithm 1). The only difference between the two stages is that the scoring stage enumerates the number of iterations that each member of the phase block spends in phase1 [1,0]. We found by ignoring several million iterative sweeps over a primary contig, the algorithm tends to be in a more favorable search space. The final phase assignment is the configuration in which each member of a phase block spent the most iterations. In practice, 50 - 100 M iterations with 10 M burn-in period generated consistent results. The limiting computational resource is memory as is not sparse.

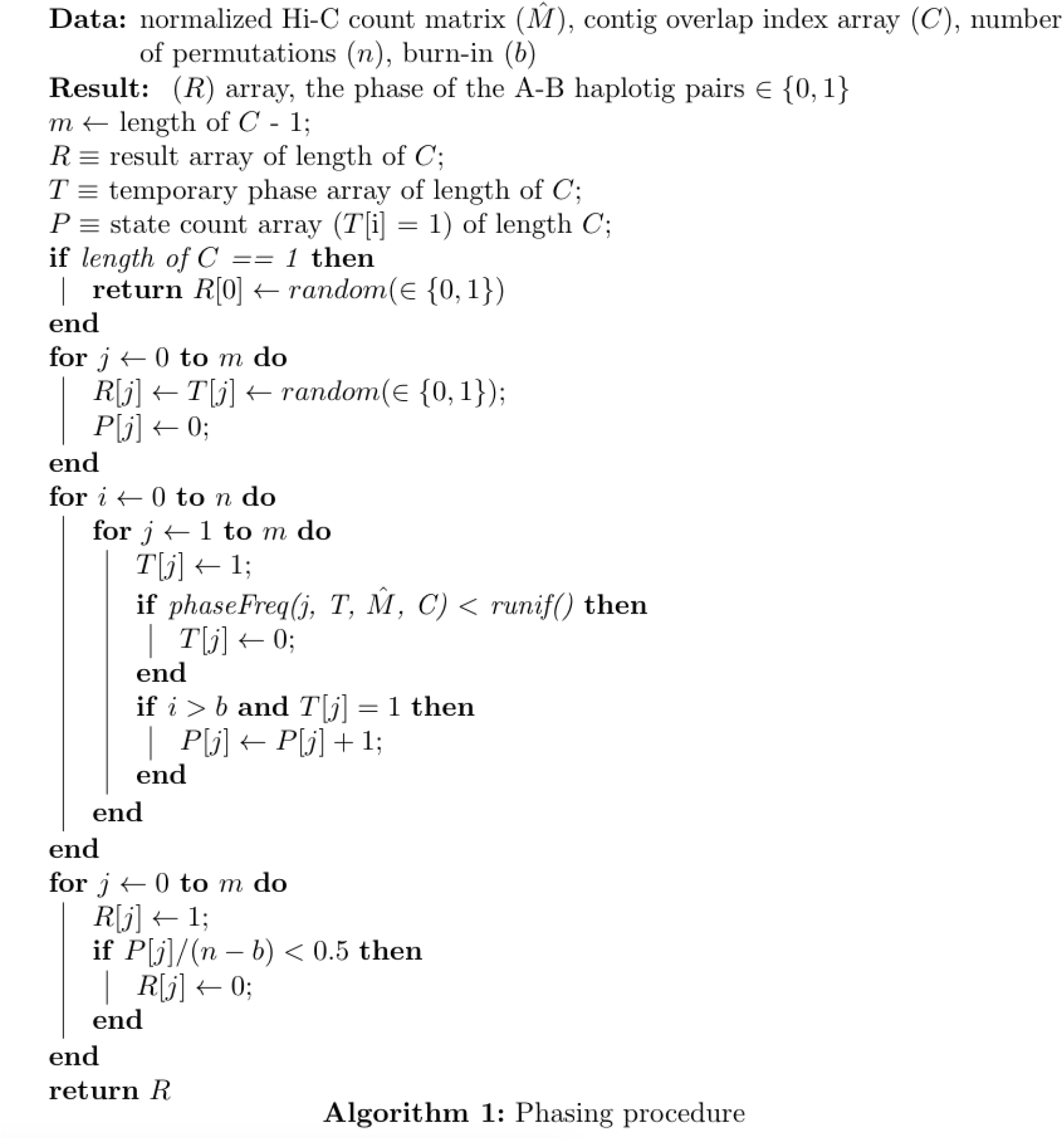

#### Emission of phased contigs

Once the phase assignments of haplotype pairs in the phase blocks are determined, the minced fasta sequences are joined into two full-length pseudo-haplotypes (phase0 and phase1) per primary contig (Fig. 1). The order of minced sequences (phase blocks plus collapsed regions) is determined by the haplotig placement file and the phase assignment is determined by the phasing algorithm. An alternate output similar to the “FALCON-Unzip” format of primary contigs and haplotigs is also available as a user-specified option.

#### Scaffolding and Scaffold-Phasing

We scaffolded the contigs from FALCON-Phase using default Proximo^20,29^ settings (Phase Genomics, WA). Briefly, reads were aligned to phase0 pseudo-haplotypes using *bwa-mem*^30^(v.0.7.15-r1144-dirty) with the -5SP and -t 8 options. *SAMBLASTER*^31^ (commit 37142b37e4f0026e1b83ca3f1545d1807ef77617) was used to flag PCR duplicates, which were later excluded from analysis. Alignments were then filtered with *samtools* (v1.5, with htslib 1.5) using the -F 2304 filtering flag to remove non-primary and supplementary alignments, as well as read pairs in which one or more mates were unmapped. Phase Genomics’ Proximo Hi-C genome scaffolding platform (commit 145c01be162be85c060c567d576bb4786496c032) was used to create chromosome-scale scaffolds from the draft assembly as previously described ^29^. As in the LACHESIS method ^20^, this process computes a contact frequency matrix from the aligned Hi-C read pairs, normalized by the number of restriction sites on each contig, and constructs scaffolds in such a way as to optimize expected contact frequency and other statistical patterns in Hi-C data. Juicebox v1.8.8 was used to correct scaffolding errors^32,33^. After scaffolding, we applied the phasing algorithm a second time, using as input the pairing of the phase0 and phase1 pseudo-haplotypes and their order along the chromosomes as determined by scaffolding.

### Method Evaluation

#### Datasets

We evaluated FALCON-Phase on three vertebrate species with different levels of heterozygosity: Puerto Rican human female, HG00733 (low), cattle (*Bos taurus indicus* x *Bos taurus taurus*, moderate), and zebra finch female (*Taeniopygia guttata*, high). For each genome, we had high coverage PacBio data for *de novo* genome assembly, Hi-C data for phasing and scaffolding, paired-end Illumina data from the parents, and trio binned Canu assemblies (Supp Table S2). The female zebra finch data was generated as part of the Vertebrate Genomes Project (Rhie et al in preparation).

#### Heterozygosity estimation

Heterozygosity was estimated two ways. First, from *k-mers* (k-length sequence) in Illumina whole genome sequencing reads (Tables S1 and S2). Fastq files were converted to fasta files, then the canonical *k-mers* were collected using meryl in canu1.7^8^ to include all the high frequency *k-mers* using the following code.

meryl -B -C -s $name.fa -m $k_size -o $name.$k

meryl -Dh -s $name.$k > $name.$k.hist

Given the *k-mer* histogram, Genomescope^34^ was used to estimate the level of heterozygosity. *k*=21 was used for human HG00733 and the cow, and *k*=31 was used for the zebra finch. A higher *k-mer* size was used for zebra finch for more accurate estimates of heterozygosity due to its higher level of polymorphism. The *k-mer* size was also used for other samples in the Vertebrate Genomes Project, from which this sample was selected.

Second, with *mummer* v 3.2.3^35^, trio binned parental Canu assemblies were aligned with *nucmer* (nucmer –l 100 -c 500 -maxmatch mom.fasta dad.fasta) and heterozygosity was computed as 1 - average identify from 1 to 1 alignments output by *dnadiff* using default parameters.

#### *De novo* genome assembly

As a precursor to FALCON-Phase, we performed *de novo* genome assembly with FALCON-Unzip^13^ using v 0.0.2 (bioconda) for zebra finch and cattle, and a binary build from August 13, 2018 for human.

Human parameters: (length_cutoff = 5000; length_cutoff_pr = 10000; pa_daligner_option = -k18 -e0.75 -l1200 -h256 -w8 -s100; ovlp_daligner_option = -k24 -e.92 -l1800 -h600 -s100; pa_HPCdaligner_option = -v -B128 -M24; ovlp_HPCdaligner_option = -v -B128 -M24; pa_HPCTANmask_option = -k18 -h480 -w8 -e.8 -s100; pa_HPCREPmask_option = -k18 -h480 -w8 -e.8 -s100; pa_DBsplit_option = -x500 -s400; ovlp_DBsplit_option = -s400; falcon_sense_option = --output-multi --min-idt 0.70 --min-cov 4 --max-n-read 200 --n-core 8; falcon_sense_skip_contained = False; overlap_filtering_setting = --max-diff 60 --max-cov 60 --min-cov 1 --n-core 12).

Bull parameters: (length_cutoff = 14850; length_cutoff_pr = 12000; pa_daligner_option = -e0.76 -l1200 -k18 -h480 -w8 -s100; ovlp_daligner_option = -k24 -h480 -e.95 -l1800 -s100; pa_HPCdaligner_option = -v -B128 -M24; ovlp_HPCdaligner_option = -v -B128 -M24; pa_HPCTANmask_option = -k18 -h480 -w8 -e.8 -s100; pa_HPCREPmask_option = -k18 -h480 -w8 -e.8 -s100; pa_DBsplit_option = -x500 -s400; ovlp_DBsplit_option = -s400; falcon_sense_option = --output_multi --min_idt 0.70 --min_cov 4 --max_n_read 200 --n_core 24; overlap_filtering_setting = --max_diff 120 --max_cov 120 --min_cov 4 --n_core 24)

Zebra finch parameters: (length_cutoff = 13653; length_cutoff_pr = 5000; pa_daligner_option = -e0.76 -l2000 -k18 -h70 -w8 -s100; ovlp_daligner_option = -k24 -h1024 -e.95 -l1800 -s100; pa_HPCdaligner_option = -v -B128 -M24; ovlp_HPCdaligner_option = -v -B128 -M24; pa_HPCTANmask_option = -k18 -h480 -w8 -e.8 -s100; pa_HPCREPmask_option = -k18 -h480 -w8 -e.8 -s100; pa_DBsplit_option = -x500 -s400; ovlp_DBsplit_option = -s400; falcon_sense_option = --output-multi --min-idt 0.70 --min-cov 2 --max-n-read 400 --n-core 24; overlap_filtering_setting = --max-diff 100 --max-cov 150 --min-cov 2 --n-core 24)

We identified and corrected chimeric contigs between non-adjacent genomic regions in the human and cattle assemblies using Juice Box Assembly Tools^33^ and D-GENIES^36^. We interrogated the contradance of the Hi-C data with the PGA scaffolds visually in JBAT. Off-diagonal signals in the heatmap are indicative of assembly/scaffolding errors. Human and cow contigs and scaffolds with discordant Hi-C signals we aligned, using *minimap2* with the -x asm5 setting, to the human or cow reference genomes (Table S2). If the contig/scaffold in question mapped chimerically (inter- or intrachromosomally) to the each species genome they were flagged. We manually broke these contigs between phase blocks and re-associated the haplotigs to the two new contigs.

To remove duplicated haplotypes in the primary contigs from the zebra finch FALCON-Unzip assembly, as suggested for highly heterozygous genomes from the Vertebrate Genomes Project (Rhie et al in preparation), we ran *purge haplotigs*^17^ on zebra finch using default settings and coverage estimates from PacBio subreads mapped to the primary contigs^17^. We recategorized 67.1 Mb of primary contigs as haplotigs (N = 632) and 25.4 Mb of repetitive sequences (N = 329) was discarded.

#### Evaluation of Phasing

Phase assignment was evaluated by *minimap2*^26^ alignments of the sequences in each phase block to the trio binned parental Canu assemblies using the -x asm5 setting. We determined the parent assignment with the higher pairwise identity of the longest alignment and required that the parental assignment of the two sequences in each phase block be concordant. The proportion of scorable length that was correctly assigned to phase was determined for each primary contig and then a weighted average was computed where the weights are the primary contig length (Table 1). By mapping each haplotig to its associated primary contig, we defined 5,273 phase blocks (haplotig plus homologous region in primary contig) in the starting FALCON-Unzip cattle assembly. 98.2% could be unambiguously assigned to a parent by FALCON-Phase; for zebra finch, 96% of the length of the 4,772 phase blocks were assigned to each parent. For human, 91.1% of the 7,774 phase blocks were assigned to each parent. In all species, the unassigned phase blocks were on average shorter than the assigned blocks (Table S4).

In addition, parent-specific *k-mers* were counted in the pseudo-haplotypes before and after contig phasing, before and after scaffold phasing, and in trio binned Canu assemblies. Parental *k-mers* were identified using Illumina data from the parents as described previously^8^ using *k*=21. Parental *k-mers* were counted in the assemblies using the “simple-dump” utility from canu v 1.7. The proportion of “correct” parental *k-mers* was used as an overall measure of contig or scaffold phasing, and was plotted for each contig or scaffold in Figure 2.

---

**Table S1.**
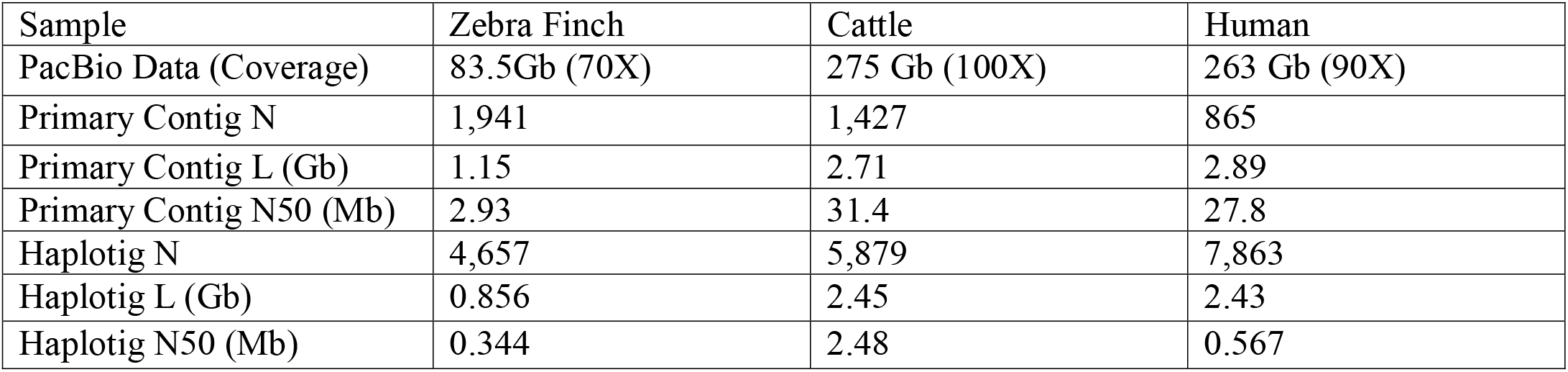
FALCON-Unzip assembly stats

**Table S2.** Data summary and availability (Genome Ark, NCBI and ENA archives)

| Sample | Zebra finch | Cattle | Human |
| --- | --- | --- | --- |
| PacBio Coverage | 73 X | 101 X | 91 X |
| PacBio Data Access | https://vgp.github.io/genomeark/Taeniopygia_guttata/ | PRJNA432857 | SRR7615963 |
| Tri Binned Parental Canu Assemblies | https://vgp.github.io/genomeark/Taeniopygia_guttata/ | GCA_003369685.2,<br>GCA_003369695.2 | https://obj.umiacs.umd.edu/marbl_publications/triobinning/h_sapiens_HG00733_dad.fasta<br>https://obj.umiacs.umd.edu/marbl_publications/triobinning/h_sapiens_HG00733_mom.fasta |
| HiC Data | 319M 2X150bp | 203M 2X80bp | 504M 2X100bp |
| HiC Data Access | VGP | PRJNA432857 | ERR1225141-ERR1225146 |
| Short Read Data | https://vgp.github.io/genomeark/Taeniopygia_guttata/ | 757M 2X150bp | PRJNA177403 |
| Short Data Access | VGP 10X | PRJNA432857 | PRJNA177403 |
| Maternal Short Read Data | https://vgp.github.io/genomeark/Taeniopygia_guttata/ | 537M 2X150bp | 710M 2X76-101bp |
| Maternal Short Read Data Access | VGP | PRJNA432857 | PRJNA42573 |
| Paternal Short Read Data | https://vgp.github.io/genomeark/Taeniopygia_guttata/ | 498M 2X150bp | 668M 2X76-101bp |
| Paternal Short Read Data Access | VGP | PRJNA432857 | PRJNA42573 |

**Table S3.** Hi-C mate pair mapping statistics

| Sample | Zebra Finch | Cattle | Human |
| --- | --- | --- | --- |
| Total Read Pairs | 625 M | 395 M | 1006 M |
| Filtered Pairs (% of total reads) | 275 M (44.1%) | 64.5M (16.3%) | 128 M (12.7%) |
| Map Distance > 10Kbp | 74.4% | 82.2% | 71.2% |
| Map Distance > 50Kbp | 59.1% | 70.6% | 50.4% |
| Map Distance > 100Kbp | 2.85% | 22.4% | 3.51% |

**Table S4.**
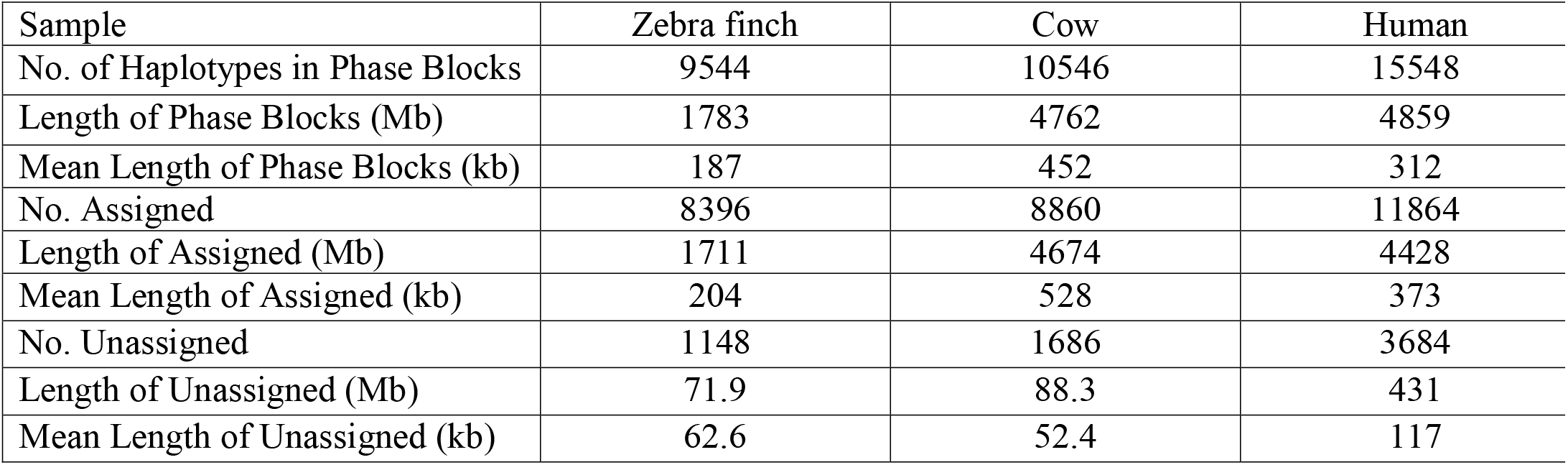
Summary statistics of parental assignment of haplotypes in phase blocks

| Sample | Zebra finch | Cow | Human |
| --- | --- | --- | --- |
| No. of Haplotypes in Phase Blocks | 9544 | 10546 | 15548 |
| Length of Phase Blocks (Mb) | 1783 | 4762 | 4859 |
| Mean Length of Phase Blocks (kb) | 187 | 452 | 312 |
| No. Assigned | 8396 | 8860 | 11864 |
| Length of Assigned (Mb) | 1711 | 4674 | 4428 |
| Mean Length of Assigned (kb) | 204 | 528 | 373 |
| No. Unassigned | 1148 | 1686 | 3684 |
| Length of Unassigned (Mb) | 71.9 | 88.3 | 431 |
| Mean Length of Unassigned (kb) | 62.6 | 52.4 | 117 |

**Table S5.** Scaffold Statistics

|  | Zebra finch | Cattle | Human |
| --- | --- | --- | --- |
| N Scaffolds | 30 | 31 | 23 |
| Total Scaffold Length (Gb) | 1.06 | 2.64 | 2.86 |
| N Contigs Scaffolded | 797 | 1040 | 514 |
| N Unscaffolded Contigs | 160 | 650 | 351 |
| Unscaffolded Contigs Length (Mb) | 7.44 | 95.1 | 38.0 |

**Figure S1.**
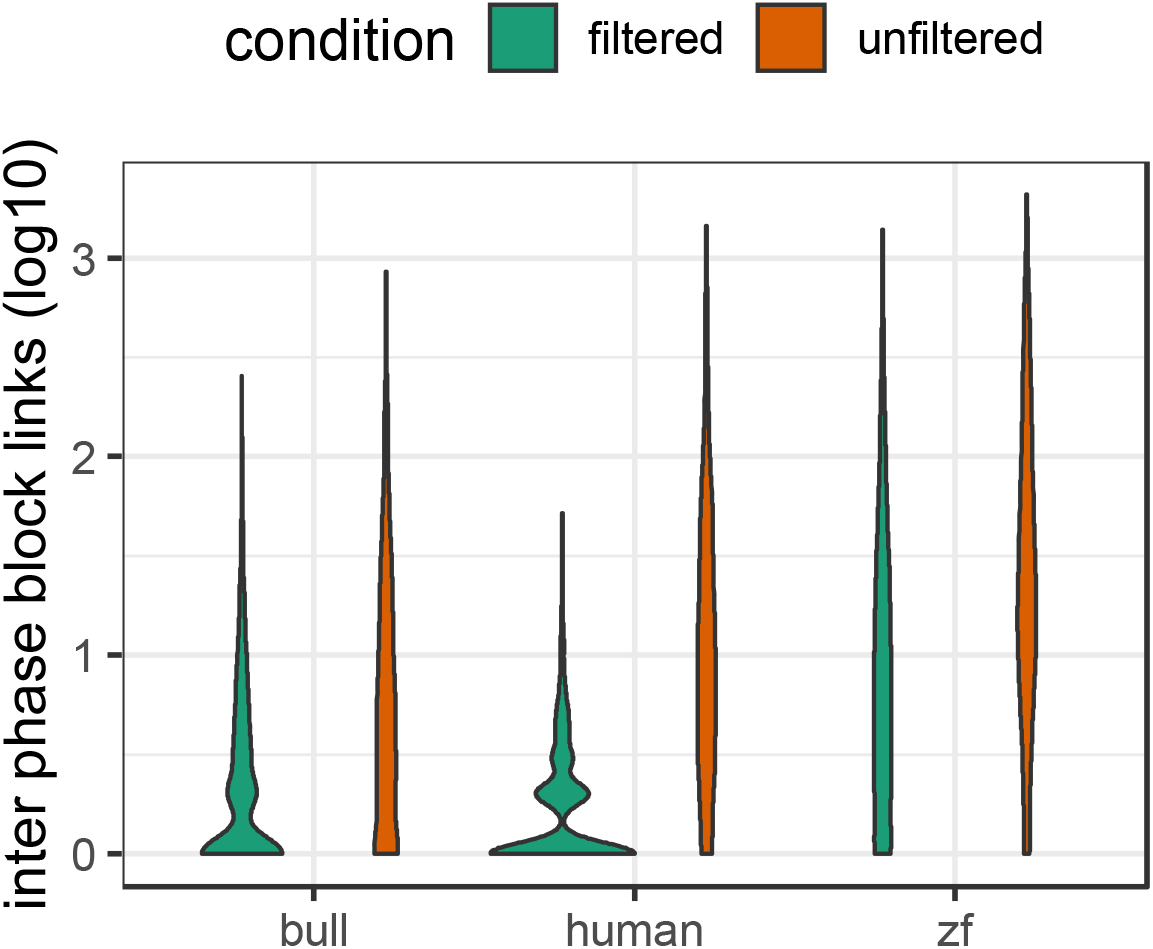
Hi-C links between phase blocks within primary contigs. Each violin plot shows the log10 distribution of Hi-C links between phase blocks before and after map quality filtering. These counts are restricted to links within primary contigs. Zero counts are not shown. The shape of this distribution is affected by an interaction between the length of long-range Hi-C contacts and the heterozygosity.

**Figure S2.**
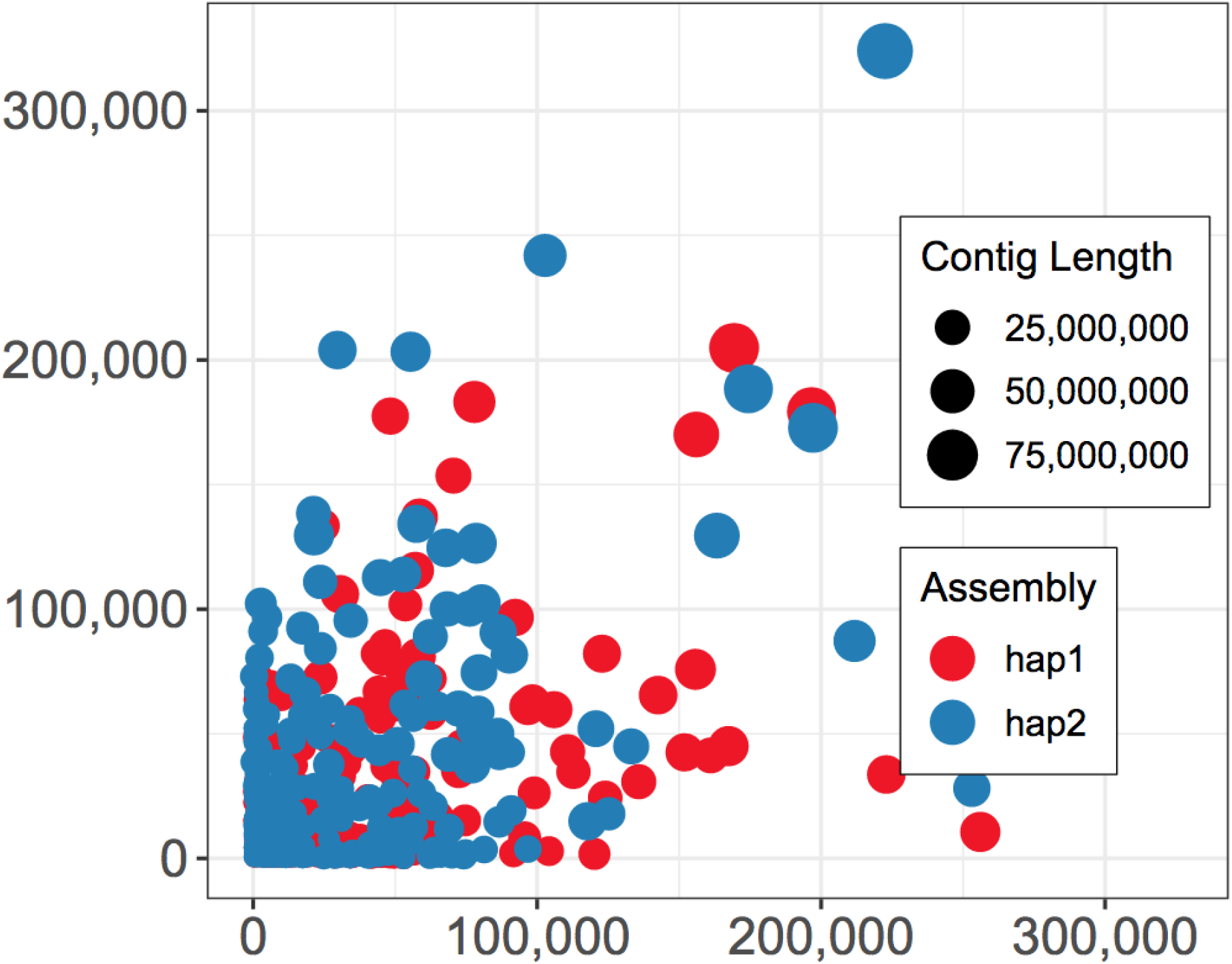
Parental *k-mer* counts for HG00733 supernova contigs (GCA_002022865.1). Markers from mother are on the X axis, father on the Y-axis. Contig or scaffold size are indicated by size of the data point.

